# Deficient mitochondrial respiration impairs sirtuin activity in dorsal root ganglia in Friedreich Ataxia mouse and cell models

**DOI:** 10.1101/2023.02.01.526688

**Authors:** Arabela Sanz-Alcázar, Elena Britti, Fabien Delaspre, Marta Medina-Carbonero, Maria Pazos-Gil, Jordi Tamarit, Joaquim Ros, Elisa Cabiscol

## Abstract

Friedreich ataxia (FA) is a rare, recessive neuro-cardiodegenerative disease caused by deficiency of the mitochondrial protein frataxin. Mitochondrial dysfunction, a reduction in the activity of iron-sulfur enzymes, iron accumulation, and increased oxidative stress have been described. However, the mechanisms causing such cellular disturbances in mammals are not completely understood. Dorsal root ganglion (DRG) sensory neurons are among the cellular types most affected in the early stages of this disease. We have previously demonstrated that frataxin depletion in primary cultures of DRG neurons results in calcium dysregulation, neurite degeneration and apoptotic cell death. However, its effect on mitochondrial function remains to be elucidated. In the present study, we found that in primary cultures of DRG neurons as well as in DRGs from the FXN^I151F^ mouse model, frataxin deficiency resulted in lower activity and levels of the electron transport complexes, mainly complexes I and II. As a consequence, the NAD^+^/NADH ratio was reduced and SirT3, a mitochondrial NAD^+^-dependent deacetylase, was impaired. We identified alpha tubulin as the major acetylated protein from DRG homogenates whose levels were increased in FXN^I151F^ mice compared to WT mice. Mitochondrial superoxide dismutase (SOD2), a SirT3 substrate, displayed increased acetylation in frataxin-deficient DRG neurons. Since SOD2 acetylation inactivates the enzyme, and higher levels of mitochondrial superoxide anion were detected, oxidative stress markers were analyzed. Elevated levels of hydroxynonenal bound to proteins and mitochondrial Fe^2+^ accumulation were detected when frataxin decreased. Honokiol, a SirT3 activator, restores mitochondrial respiration. Altogether, these results provide the molecular bases to understand mitochondria dysfunction in sensory neurons which have greater susceptibility to frataxin deficiency compared to other tissues.

## 1. Introduction

Frataxin is a mitochondrial protein that, at concentrations below 40% of normal levels, leads to Friedreich ataxia (FA), a disease caused by the presence of large expansions of GAA triplet repeats in the first intron of the frataxin (FXN) gene, which results in a low transcription rate and, consequently, diminished frataxin expression [1]. While triplet expansions in homozygosis are the most observed genetic abnormality in FA patients, 4% of them are compound heterozygotes displaying GAA repeats in one allele and point mutations in the other [2]. Decreased expression of frataxin is associated with mitochondrial dysfunction, iron and calcium imbalance, and increased oxidative stress. The function of frataxin and the mechanisms causing such cellular disturbances are not completely understood but it has been shown to regulate the activity of cysteine desulfurase, an enzyme required for the biosynthesis of iron–sulfur clusters [3]. Although it is generally accepted that frataxin activates iron–sulfur biogenesis in eukaryotes, iron– sulfur deficiency is not a universal consequence of frataxin deficiency, suggesting that the role of frataxin in this process is not essential [4].

Patients with FA suffer progressive limb and gait ataxia, dysarthria, reduced tendon reflex, extensor plantar responses and loss of position and vibration senses. Although much research has been conducted [5], there is no effective cure for the disease. The pathologic changes occur first in dorsal root ganglia (DRG) with loss of large sensory neurons, followed by degeneration of the spinocerebellar and corticospinal tracts [6]. DRG neurons express the highest levels of frataxin and display high vulnerability to frataxin down-regulation. For this reason, the consequences of frataxin depletion have been studied, at the histological level, in DRGs using conditional knockout mice [7] and samples from patients with FA [8] [9]. Also, using primary cultures of frataxin-deficient DRG neurons, our group observed alterations of several parameters such as a decrease in mitochondrial membrane potential, increased fodrin cleavage by calpain and caspase, and Bax induction [10] [11]. These events led to apoptotic cell death that was rescued either by supplementing cultures with calcium chelators such as BAPTA or with TAT-BH4, the antiapoptotic domain of Bcl-xL fused to TAT peptide [10]. Besides apoptosis, ferroptosis is a process of cell death involving iron accumulation, lipid peroxidation and decreased GPX4 activity, among other factors [12] that has also been described in several models of FA [13] [14].

Our group has previously reported that, in frataxin-deficient cardiomyocytes and in DRG neurons, opening of the mitochondrial permeability transition pore occurred [15] and can be triggered by, among other causes, increased acetylation of the mitochondrial cyclophilin D (CypD) due to impairment of Sirt3 function, the most abundant mitochondrial sirtuin [16] [17]. Sirt3 belongs to the sirtuin family, which are NAD-dependent class III histone deacetylases (HDAC). It is synthesized as a 44-kDa precursor that, once processed by mitochondrial proteases, is converted to a mature form of 28 kDa [18]. Although Sirt4 and Sirt5 have also been found in mitochondria [19], their deacetylase activity is very weak compared to that of SirT3 [20]. For this reason, SirT3 is the main sirtuin responsible for controlling the lysine acetylation levels of mitochondrial proteins [21]. In this context, SirT3 has been involved in the control of mitochondrial fatty acid oxidation, and regulates the activity of succinate dehydrogenases, pyruvate dehydrogenase as well as tricarboxylic acid cycle enzymes such as aconitase, isocitrate dehydrogenase and malate dehydrogenase [22] [23] [24]. In the electron transport chain (ETC), several proteins from complexes I to V are SirT3 targets [19] [23]. Protein acetylation has been observed in the heart of a cardiac mouse model of FA and increases with age. Acetyl-SOD2 was identified as one of the targets; the acetylated form is inactive and, as a result, ROS production increases. Higher acetylation was associated with impaired mitochondrial fitness, altered lipid metabolism, and a decline in heart function as well as the development of steatosis and cardiac fibrosis [25]. Sirt3 activators have thus been tested as a possible strategic therapy. Among them, resveratrol (an SirT3 activator [26] that increases SirT3 levels) was suggested as a possible therapy for FA a few years ago [5] and is now in phase 2 trials. Honokiol, another SirT3 activator, has anti-inflammatory, anti-tumor, anti-oxidative, and neuroprotective properties [27]. It has proven to be effective at, among other functions, reducing SOD2 acetylation, thus maintaining SOD2 activity under doxorubicin-induced oxidative stress [28].

In the current study, we analyzed the consequences of frataxin deficiency in DRG neurons using two models: primary cultures of frataxin-deficient DRGs and DRGs from the new mouse model FXN^I151F^ [29]. This mouse carries the I151F point mutation, equivalent to the human I154F pathological mutation. FXN^I151F^ homozygous mice present very low frataxin levels, biochemical alterations, and neurological deficits (starting at 23 weeks of age) that mimic those observed in patients with FA. In summary, we detected a deficiency of mitochondrial respiration, which resulted in a decreased NAD^+^/NADH ratio and, as a consequence, a decline in sirtuin activity. Mitochondrial SOD2, a SirT3 substrate, showed increased levels of its acetylated (inactive) form, leading to oxidative stress. Interestingly, restored mitochondrial function was observed in cells supplemented with honokiol, a SirT3 activator. These results indicate that in frataxin-deficient DRGs a negative feedback mechanism involving decreased SirT3 activity contributes to neuron lethality.

## 2. Materials and Methods

### 2.1. Animals and isolation of DRGs from FXN^I151F^ mouse model

All experimentation with animals was performed according to the National Guidelines for the regulation of the use of experimental laboratory animals issued by the Generalitat de Catalunya and the Government of Spain (article 33.a 214/1997), which comply with the ARRIVE guidelines. Experimental protocols were evaluated and approved by the Experimental Animal Ethical Committee of the University of Lleida (CEEA). All procedures were performed at the animal facility. For euthanasia of animals, the guidelines of the American Veterinary Medical Association (AVMA) were followed. Male and female Sprague-Dawley rats (RRID:RGD_70508) were maintained in standard conditions of 12-hour cycles of light and dark, a constant temperature of 20°C and eating and drinking ad libitum.

FXN^I151F^ heterozygous mice (C57BL/6J-Fxnem10(T146T,I151F)Lutzy/J) were obtained from the Jackson Laboratory, Bar Harbor, ME, USA (Stock Number 31922) as previously described [29]. Intercrosses of heterozygous animals were performed to generate the homozygous WT and FXN^I151F^ mice. Animals were housed in standard ventilated cages with 12 h light/dark cycles and fed with a normal chow diet ad libitum. Animals were weighed weekly. Genotyping was performed by sequencing a PCR product amplified from DNA extracted from tail biopsy specimens as described previously [30]. For DRG isolation, animals were sacrificed by cervical dislocation at 21 or 39 weeks of age, dissected and snap-frozen immediately in liquid nitrogen, and stored at -80°C.

### 2.2. Isolation and culture of primary rat DRG sensory neurons

For primary culture of DRG neurons, DRG were extracted from neonatal Sprague–Dawley rats (P3–P4) and purified as described [10] [11] with modifications. The ganglia (without the nerve roots) were dissociated by incubation in 0.025% trypsin (Sigma-Aldrich) without EDTA in GHEBS (137 mM NaCl, 2.6 mM KCl, 25 mM glucose, 25 mM HEPES, 100 μg/mL penicillin/streptomycin) for 30 min. Ganglia were then gently disrupted using a pipette to obtain a single cell suspension in Neurobasal culture media (GIBCO, Cat# 21103049) enriched with 2% horse serum (GIBCO, Cat# 16050-122), 2% B27 Supplement (ThermoFisher Scientific, Cat#17504-044), 0.5 mM L-glutamine (GIBCO, Cat# 25030-024), 100 U/mL penicillin plus 100 ng/mL Streptomycin (GIBCO, Cat# P4458). Aphidicolin (Sigma–Aldrich, Cat# A0781) was added at a final concentration of 3.4 mg/mL to prevent the growth of non-neuronal cells and supplemented with 50 ng/mL murine β-nerve growth factor (PeproTech, Cat# 450-34) with 3 mg/mL DNAse I grade II (Roche, Cat# 104159). The cell suspension was centrifuged at 1,300 rpm for 5 minutes with 7.5% BSA solution (Sigma, Cat# A8412). After 1 h of pre-plating in a p60 tissue dish (Corning Incorporated, Cat# 35004) at 37°C/5%CO_2_, the cells were then plated in tissue dishes pre-treated with 0.1 mg/mL of collagen (Sigma, Cat# C7661-25) at a cell density of 14,000 cells/well. After 1–2 days, lentivirus transduction was performed with shRNA sequences targeting frataxin mRNA as described previously [31]. The RefSeq used was NM-008044, which corresponds to mouse frataxin. The clones used were TRCN0000197534 and TRCN0000006137 (here referred to as FXN1, FXN2). The vector SHC002, a non-targeted scrambled sequence, served as a control (Scr). Lentivirus particles (20 ng/1,000 cells) were added and replaced with fresh medium 6 hours later. Experiments were performed after 1, 3 or 5 days as indicated.

### 2.3. Mitochondrial respiration in primary cultures of DRG neurons

Oxygen consumption rate (OCR) and extracellular acidification rate (ECAR) were measured in primary cultures of DRG neurons using the Seahorse XFp analyzer (Agilent). DRG neurons were cultured (1, 3 and 5 days after lentivirus transfection) in Seahorse XF cell culture plates (103726-100) pre-treated with collagen. One hour before the assay, the culture medium was changed to DMEM (Dulbecco’s modified Eagle’s medium, pH 7.4) with 2 mM glutamine, 1 mM pyruvate, and 10 mM glucose, the plates were placed in a 37ºC non-CO_2_ incubator, and OCR and ECAR were measured according to the manufacturer’s instructions. Mitochondrial complex inhibitors were sequentially injected into each well: oligomycin (1.5 µM) to inhibit complex V, FCCP (3 µM) as an uncoupling agent, and rotenone-antimycin A (0.5 µM each) to inhibit complexes I and III. All data were automatically calculated, recorded, and plotted using the Seahorse analytics software, and normalized by protein concentration per well. To study the role of SirT3, 2 µM honokiol was added to primary cultures 0 and 2 days after lentivirus transduction, and cellular respiration was analyzed using Seahorse XFp analyzer 48 h later.

### 2.4. Enzymatic activity of mitochondrial respiratory complexes

DRGs isolated from WT and FXN^I151F^ mice were homogenized with a Dounce homogenizer (ten strokes) in 50 mM Tris-HCl, 0.15 M KCl, pH 7.4 and gently sonicated. The homogenate was centrifuged (800 *x g* for 15 min) and the supernatants were used to assay CoQ-dependent respiratory chain activities (CI + III and CII + III) as previously described [32] with minor modifications. To assess complex I + III activity, the rate of cytochrome c reduction was measured at 550 nm, using NADH as the electron donor. The sample was incubated in 50 mM Tris-HCl, 0.15 M KCl, pH 7.4 containing 1 mg/mL BSA, 40 µM cytochrome c and 240 µM KCN. After 2–3 min, 0.8 mM NADH was added, and cytochrome c reduction was recorded for another 2–3 minutes. The same procedure was performed in the presence of 4 µM rotenone (a complex I inhibitor) to determine the rotenone-sensitive reduction of cytochrome c. To check the activity of complex II + III, the sample was incubated for 10 minutes with 50 mM Tris-HCl, 0.15 M KCl, pH 7.4 plus 2 mM EDTA, 1 mg/mL BSA, 240 µM KCN, 4 µM rotenone and 10 mM succinate as a substrate. The reaction was initiated by addition of 40 µM cytochrome c and the decrease in absorbance was monitored at 550 nm for 2–3 minutes. The assay was also performed in the presence of 0.5 mM 2-thenoyltrifluoroacetone, a complex II inhibitor. In both assays (complex I + III and complex II + III), the results were expressed in nmol reduced cyt c/min/mg prot. Citrate synthase activity was measured with a coupled assay to reduce 5,5′-dithiobis-(2-nitrobenzoic acid) (DTNB) [33]. Briefly, DRG extracts obtained as described above were centrifuged at 12,000 rpm for 10 min and supernatants were added to 100 mM Tris-HCl pH 8.1 with 0.4 mg/mL DTNB and 10 mg/mL acetyl-CoA. Absorbance was measured at 412 nm for 2 min. Then, 8.5 mg/mL of oxaloacetate were added to the cuvette and the absorbance was measured again at 412 nm for 2 min for the detection of reduced DTNB. All spectrophotometric measurements were performed in 0.1-mL cuvettes using a double beam spectrophotometer (Shimadzu UV-160).

### 2.5. Western blot analysis

To obtain homogenates from primary cultures, DRG neurons were lysed (at day 5 after lentivirus transduction) with 2% SDS, 125 mM Tris-HCl pH 7.4, plus protease inhibitor cocktail (Roche), phosphatase inhibitor (Roche), and 5 µM trichostatin A (TSA, Sigma). Cell extracts were sonicated, heated at 95ºC for 5 min and centrifuged at 10,000 rpm for 10 min. Protein concentration was measured using the microBCA method (Thermo Scientific). To obtain tissue homogenates, isolated DRGs were placed in 1.5 mL screw cap polypropylene tubes in the presence of lysis buffer consisting of 50 mM Tris-HCl pH 7.5 containing a protease inhibitor cocktail (Roche). Glass beads (0.5–1.0 mm) were added to the mixture, which was then homogenized in a BioSpec Mini-Beadbeater. Following homogenization, SDS was added to a final concentration of 4%. This homogenate was vortexed for 1 min, heated at 98ºC for 5 min, sonicated and subsequently centrifuged at 12,000 *x g* for 10 min. Protein content in the supernatant was quantified using the BCA assay (Thermo Scientific).

Protein extracts (10–15 µg) were subjected to SDS-polyacrylamide gel electrophoresis and transferred to PVDF membranes or nitrocellulose membranes and blocked with I-Block (ThermoFisher, T2015). Primary antibodies used were: frataxin (1:1000 Abcam, ref. Ab219414), OxPhos (1:20,000 Invitrogen, ref. 458,099), SOD2 (1:2,000 AbCam, ref. Ab13533), acetylated Lys68 SOD2 (1:2,000 AbCam, ref. Ab137037), acetylated Lys40 alpha tubulin (1:20,000 Proteintech, ref. 66200), SirT3 (1:2,000 Proteintech, ref. 10099-1AP to detect mouse SirT3), SirT3 (1:1,000 Invitrogen, ref. MAB14910 to detect rat SirT3), beta-actin (1:50,000 Chemicon, ref MAB1501R), 4-Hydroxynonenal (HNE)-Michael adducts (1:2,000 Calbiochem, ref. 393207) and acetylated-lysine (1:2,000 Cell Signaling, ref. CST9681S). Membranes were stained with Coomassie brilliant blue for normalization.

### 2.6. NAD^+^ and NADH measurement

NAD^+^ and NADH measurement in DRG tissue or cell cultures was carried out using the EnzyChromTM NAD^+^/NADH Assay Kit (E2ND-100) according to the manufacturer’s protocol. In brief, DRG tissue or cell cultures were homogenized with 100 μL of NAD^+^ or NADH extraction buffer. Extracts were heated for 5 minutes at 60ºC and 20 µL of the assay buffer was added, followed by 100 μL of the opposite buffer extraction (to neutralize the extracts). Mixtures were centrifuged at 14,000 rpm for 5 minutes and 40 μL of the supernatant were transferred into a 96-well plate and assayed as indicated. Optical density at 565 nm was recorded at time zero, and after 30 minutes, using a 96-well plate reader spectrophotometer. A standard curve was performed to determine the NAD^+^/NADH concentration of the samples.

### 2.7. Sirtuin activity

Sirtuin activity was measured using a SirT3 activity assay kit (Abcam, ab156067) in line with the manufacturer’s protocol. DRGs or cultured cells were resuspended in PBS, sonicated, and centrifuged at 14,000 rpm for 5 min. Trichostatin A, which selectively inhibits the class I and II mammalian histone deacetylases, but not class III (sirtuins) was added to give a final concentration of 5 µM. The fluorescence intensity was measured for 30 min at 5 min intervals on a fluorometric microplate reader (Tecan infinity M200 pro) with excitation at 340–360 nm and emission at 440–460 nm. Data were normalized by protein concentration. Sirtuin activity was represented as relative levels of fluorescence intensity.

### 2.8. Two-dimensional (2D) electrophoresis

Isolated DRGs from mice were disrupted under liquid nitrogen using a mortar and pestle, and ground to a fine powder. The powder was resuspended in rehydration buffer containing 7 M urea, 2 M thiourea, 4% Chaps, 50 mM dithiothreitol (DTT) plus protease inhibitors (Roche). After incubation (30 min in a shaker at 25ºC), 0.5% (v/v) ampholytes, pH 3–10 (GE Healthcare Life Sciences), and bromophenol blue were added and centrifuged (14,000 rpm for 10 min) to remove insolubilized material. Isoelectric focusing was performed in immobilized pH gradient strips (17 cm, pH 3–10 NL; Bio-Rad). The focused strips were stored at −80°C until second-dimension electrophoresis was performed. Thawed strips were equilibrated for 15 min in 370 mM Tris-HCl pH 8.8 containing 6 M urea, 2% (w/v) SDS, 20% (v/v) glycerol, and 130 mM DTT and then reequilibrated for 15 min in the same buffer containing 135 mM iodoacetamide in place of DTT. Second-dimension SDS-PAGE was performed on 11% polyacrylamide gels. Gels were stained with the fluorophore Oriole (Rio-Rad) or transferred to PVDF for western blot anti acetyl-Lys. In both cases, images were scanned with a Chemidoc CCD camera (Bio-Rad) and analyzed with PDQuest software (Bio-Rad). Acetylated protein was excised from the Oriole stained gel, and identified after in-gel trypsin digestion followed by mass spectrometry at the Mass Spectrometry and Proteomics Core Facility of the Institut de Recerca Biomèdica de Barcelona (IRBB).

### 2.9. Mitochondrial superoxide and mitochondrial iron

Mitochondrial superoxide content was detected with the MitoSOX Red probe (ThermoFisher, M36008), and mitochondrial Fe^2+^ content was determined with the Mito-FerroGreen probe (Dojindo, M489-10) according to the manufacturer’s instructions. In both cases, DRG neurons were placed in ibiTreat 4-well m-slides (Ibidi, Cat#80286) pre-treated with 0.1 mg/mL of collagen. Three days after lentivirus transduction, cells were washed three times with warm buffer (Hank’s buffered saline solution and 10 mM HEPES medium) and incubated for 10 min with 5 µM MitoSox probe (510/580 nm), or for 30 min with 5 µM Mito-FerroGreen probe (488/500 nm) diluted in Hank’s buffered saline solution medium. Cells were washed three times and images recorded by confocal microscopy (Olympus FV1000) with a 100x objective. Mitochondrial fluorescence intensity per cellular soma was quantified using Image J analysis software.

### 2.10. Statistical analysis

Data are presented as means ± SD. Statistical analysis was performed using a two-tailed Student t-test. P-values lower than 0.05 (*), 0.01 (**) or 0.005 (***) were considered significant. GraphPad Prism 5.0 was used for all analyses and graphs. For primary cultures of DRG neurons, the data were obtained from at least three independent isolations (referred to as n number). For each isolation, an average of 15 rats were used.

## 3. Results

### 3.1. Mitochondrial impairment in primary cultures of frataxin-deficient DRG neurons

Several reports have demonstrated that mitochondrial respiration is impaired in different tissues as a consequence of decreased frataxin levels. Among them, heart has been the most studied [34] [35], but it has also been demonstrated in other FA models, like skin fibroblasts, lymphoblasts, and lymphocytes [36] [37]. However, how frataxin affects sensory neurons, the primary target in FA, has remained unresolved [38]. In this study, to determine whether mitochondrial respiration was also affected in DRG neurons, they were isolated from neonatal rats and transduced with lentiviral vectors carrying frataxin-interfering shRNA (FXN1 or FXN2) or a non-interfering shRNA (Scr) as a control. Using this model, we observed that 5 days after transduction, frataxin levels were reduced to 20–30%, the mean values for shRNAs FXN1 and FXN2 respectively (Supplementary Fig. 1), which were similar to those values found in FA patients. However, as we have previously shown, as early as 24 h after transduction, frataxin levels began to decline, and this decline continued for 5 days after transduction with frataxin targeting shRNAs [11]. The seahorse technology allowed us to determine the time course effect of decreased frataxin levels on mitochondrial respiration in primary cultures of DRG neurons. The oxygen consumption rate (OCR) was measured under several conditions: basal respiration, oligomycin addition (a complex V inhibitor), FCCP addition (an uncoupling agent) and rotenone/antimycin A addition (rotenone inhibits complex I and antimycin A inhibits complex III). As shown in Fig. 1A, the decrease in OCR was correlated with frataxin decay. Using this data, several parameters were quantified 5 days after transduction (Fig. 1B). Basal respiration, ATP-linked respiration, maximal respiration and spare capacity, all decreased following transduction with shRNAs FXN1 and FXN2 compared to non-interfering shRNA (Scr). In addition, the mitochondrial ATP production rate (mitoATP) versus cytosolic ATP production rate (GlycoATP) was determined by combining OCR data with extracellular acidification rate (ECAR) (Fig. 1C). As shown in Fig. 1D, FXN1 and FXN2 cells displayed a huge reduction (80–85%) in mitoATP production rate compared to Scr cells. These results are consistent with previous results from our group demonstrating that frataxin reduction induced mitochondrial depolarization in primary cultures of DRG neurons (Supplementary Fig. 2).

**Fig. 1.**
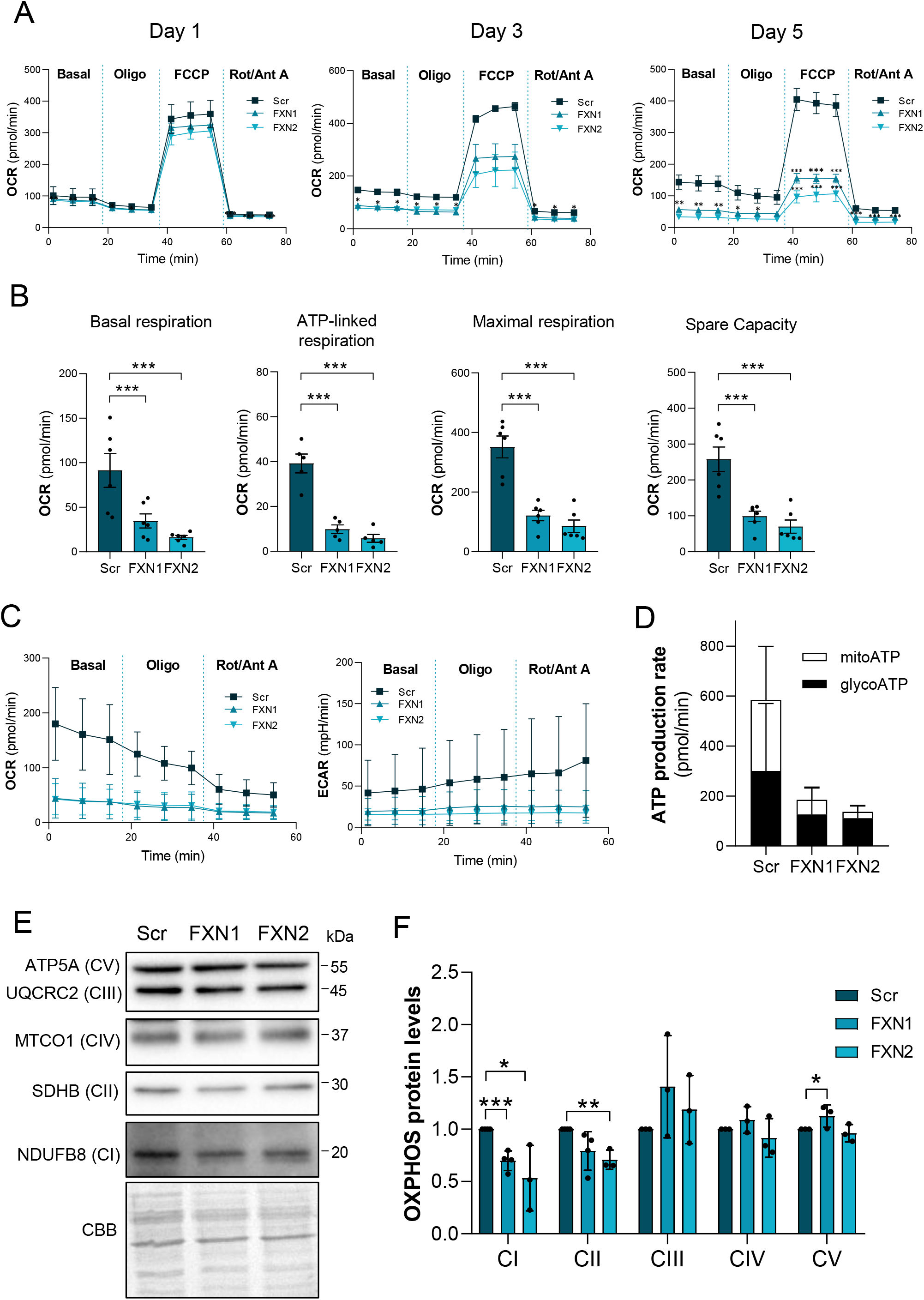
Mitochondrial ETC complexes are impaired in primary cultures of frataxin-deficient DRG neurons. (A) Mitochondrial stress analysis of frataxin-deficient DRG neurons (FXN1 and FXN2) compared to control cells (Src) at day 1, 3 and 5 after lentivirus transduction (OCR: Oxygen consumption rate) (n=2 independent cultures at day 1 and 3; n=6 independent cultures at day 5). (B) Basal respiration, ATP production, maximal respiration rate, and spare capacity of FXN1 and FXN2 neurons compared to Src at day 5 after lentivirus transduction (n=6 independent cultures). (C) OCR and extracellular acidification rate (ECAR) of FXN1 and FXN2 cells compared to control Scr cells at day 5 after lentivirus transduction (n=3 independent cultures). (D) Mitochondrial and glycolytic ATP production rates were obtained from C. (E) Indicated proteins of the mitochondrial OXPHOS system were analyzed by western blot in Scr, FXN1 and FXN2 primary culture homogenates (at day 5 after lentivirus transduction). Representative western blot images are shown. (F) Histograms represent the mean ± SD from (n=3 independent experiments). Coomassie Brilliant Blue (CBB) protein stain was used as a loading control. Data are the mean ± SD. Significant differences between Scr and FXN1 or FXN2 are indicated (p values < 0.05(*), 0.01(**), or 0.001(***)).

Since frataxin is involved in Fe-S cluster synthesis, it is possible that such deficiencies in activity are due to decreased protein levels, as several proteins in electron transport chain (ETC) complexes contain Fe-S. To this end, mitochondrial ETC complexes were analyzed by western blot from primary cultures of DRG neurons (Fig. 1E). Cell extracts of FXN1 cultures displayed around a 50% reduction in SDHB (complex II) and NDUFB8 (complex I) protein levels compared to Scr cultures (Fig. 2B). Nevertheless, no differences in UQCRC2 (complex III), MTCO1 (complex IV) or ATP5A (complex V) were found between FXN1 and Scr cultures (Fig. 1F).

**Fig. 2.**
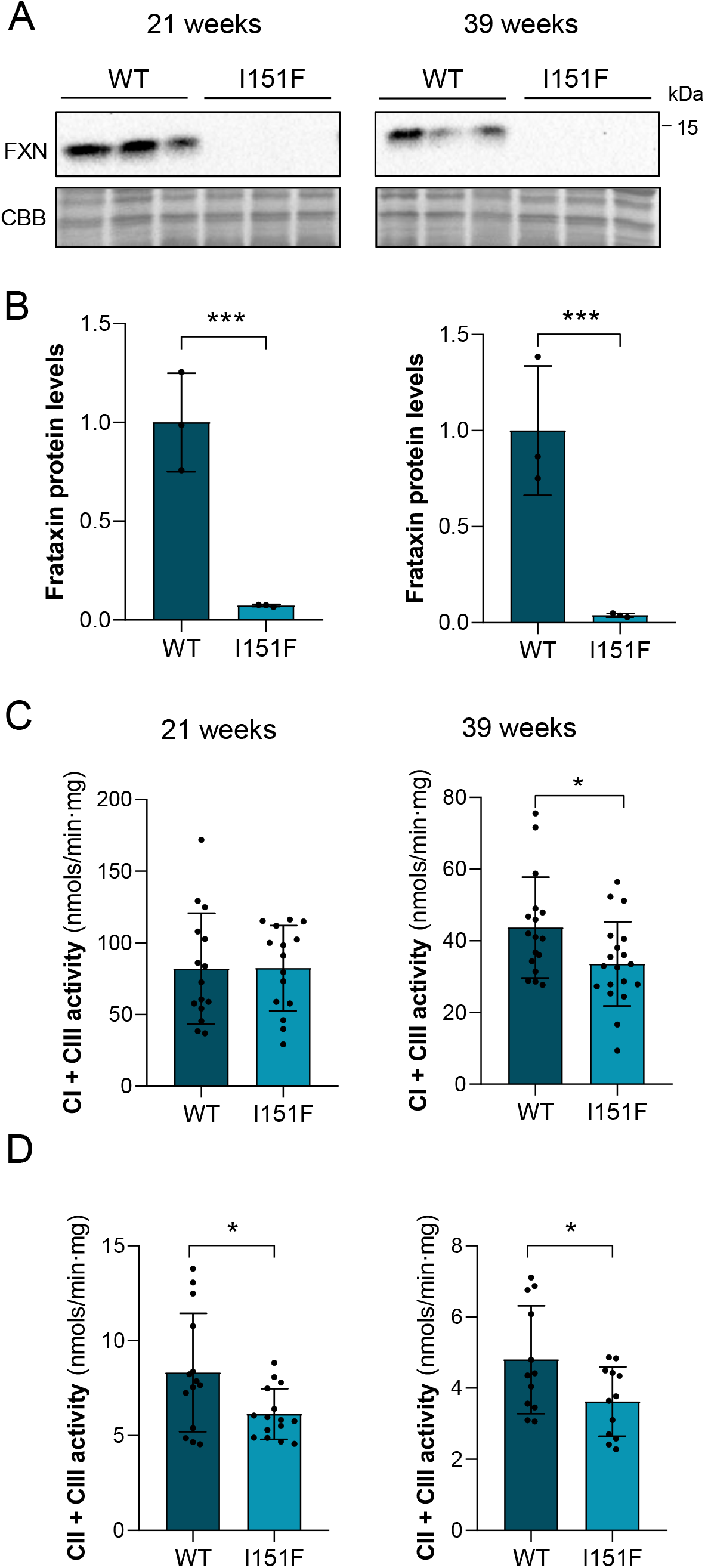
DRGs from FXN^I151F^ mice present reduced mitochondrial complex I and II activities. (A) Relative mature frataxin (15 kDa) content analyzed by western blot in DRG homogenates from 21- and 39-week-old WT and FXN^I151F^ mice. Representative images from frataxin western blot membranes under nonsaturating conditions are shown. (B) Data represent the mean ± SD from three different mice. Coomassie Brilliant Blue (CBB) stain was used as a loading control. (C) Complex I+III, and (D) complex II+III activities were measured in DRG homogenates from 21- and 39-week-old WT and FXN^I151F^ mice. Data are the mean ± SD from n=3–4 animals/group. Significant differences between WT and FXN^I151F^ mice are indicated (p values < 0.05(*), 0.01(**) or 0.001(***)).

### 3.2. Mitochondrial complex I and II are impaired in DRGs from FXN^I151F^ mouse model

To analyze whether such differences also occur *in vivo*, we used the new mouse model FXN^I151F^ carrying a missense point mutation equivalent to human I154F [29]. In homozygosity, these mice present low levels of frataxin in all tissues and start to show neurological symptoms at 23 weeks of age. It is known that they present mitochondrial alterations that are more marked in the nervous system (cerebrum and cerebellum) than in the heart, but DRG neurons have not been analyzed previously [29].

Here, we isolated DRGs from 21- and 39-week-old WT and FXN^I151F^ mice. Frataxin levels in the mutant mice were 3–8% compared to WT mice both at 21 and 39 weeks old (Fig. 2A and B). To study mitochondrial function, the enzymatic activity of mitochondrial ETC complexes was determined. As shown in Fig. 2C, at 21 weeks of age, DRG homogenates from FXN^I151F^ mice showed no differences in complex I + III activity compared to WT mice. However, differences were statistically significant at 39 weeks. Interestingly, complex II + III activity showed a significant reduction in FXN^1515F^ mice compared to WT at 21 weeks, as well as at 39 weeks (Fig. 3D). Citrate synthase has usually been used as an indication of mitochondrial content. Thus, we measured citrate synthase activity, and no differences were observed (Supplementary Fig. 3). Next, the levels of several proteins from the mitochondrial ETC from DRG homogenates from 21- and 39-week-old WT and FXN^I151F^ mice were analyzed (Fig. 3A). At 21 weeks of age, only mitochondrial SDHB from complex II showed significant differences (Fig. 3B). At 39 weeks of age, there was a decrease in NDUB8 from complex I and in SDHB from complex II. At both ages, no differences were found in UQCRC2 (complex III), MTCO1 (complex IV) or ATP5A (complex V) in FXN^151F^ versus WT animals (Fig. 3B).

**Fig. 3.**
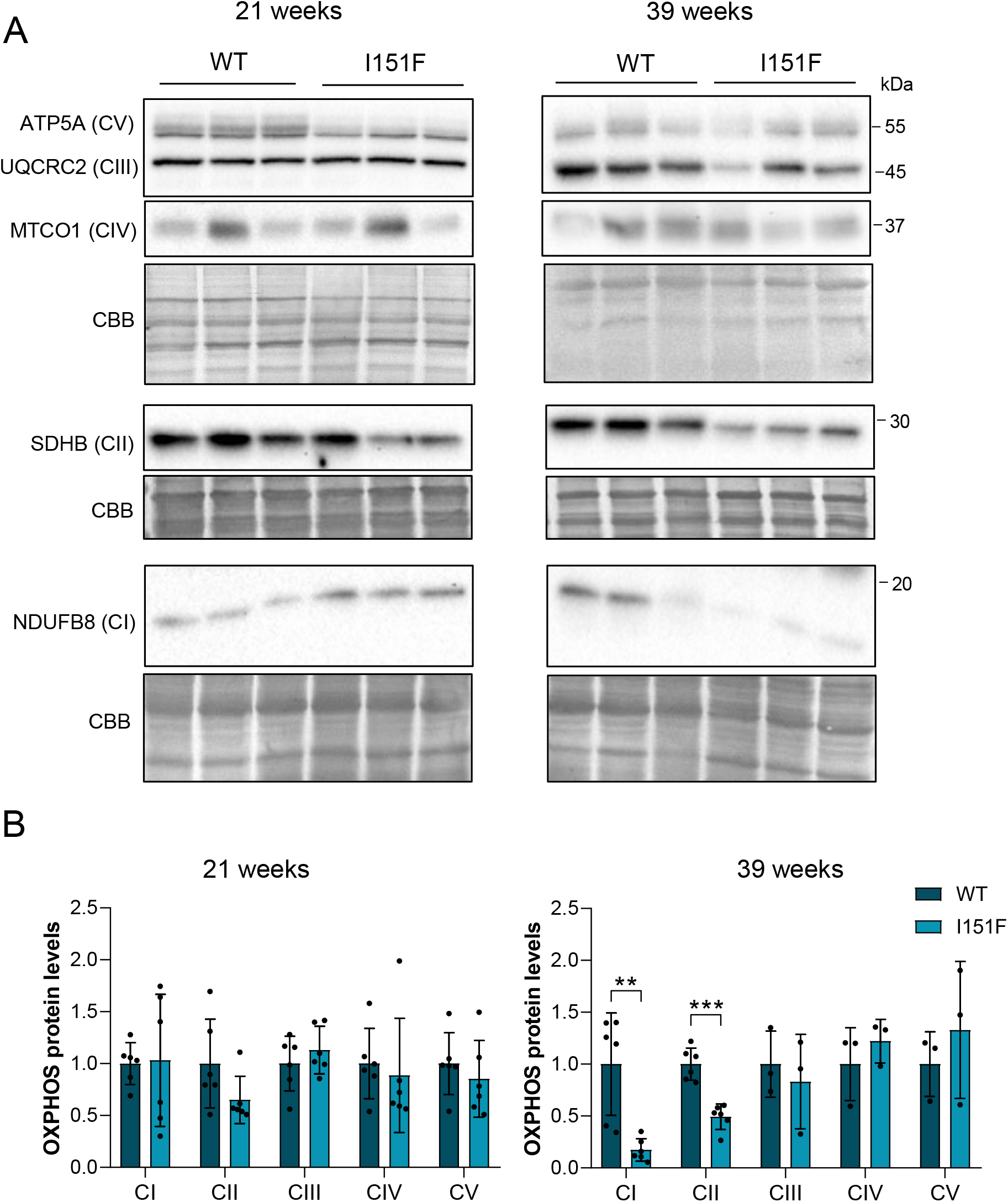
Western blot analysis of components of the OXPHOS system. Indicated proteins of the mitochondrial OXPHOS system were analyzed by western blot of DRG homogenates from 21- and 39-week-old WT and FXN^I151F^ mice. (A) Western blot images are shown. (B) Histograms represent the mean ± SD from n=3–6 animals/group. Coomassie Brilliant Blue (CBB) protein stain was used as a loading control. Significant differences between Scr and FXN1 or FXN2 are indicated (p values < 0.05(*), 0.01(**), or 0.001(***)).

### 3.3. Reduced NAD^+^/NADH ratio and decreased sirtuin activity in frataxin-deficient DRG cells

The results described above showed that deficiency of frataxin induced an impairment in the mitochondrial ETC, mainly affecting complex I and II. These alterations might lead, in addition to an ATP deficiency (Fig. 1D), to other consequences such as alterations in the NAD^+^/NADH ratio. As shown in Fig. 4A, this ratio was highly reduced (5-fold) in frataxin-deficient primary cultures of DRG neurons (FXN1) compared to control cultures (Src). Although differences were not as great, a decreased NAD^+^/NADH ratio was also found in DRG homogenates from both 21- and 39-week-old mutant mice compared to WT mice (Fig. 4B).

**Fig. 4.**
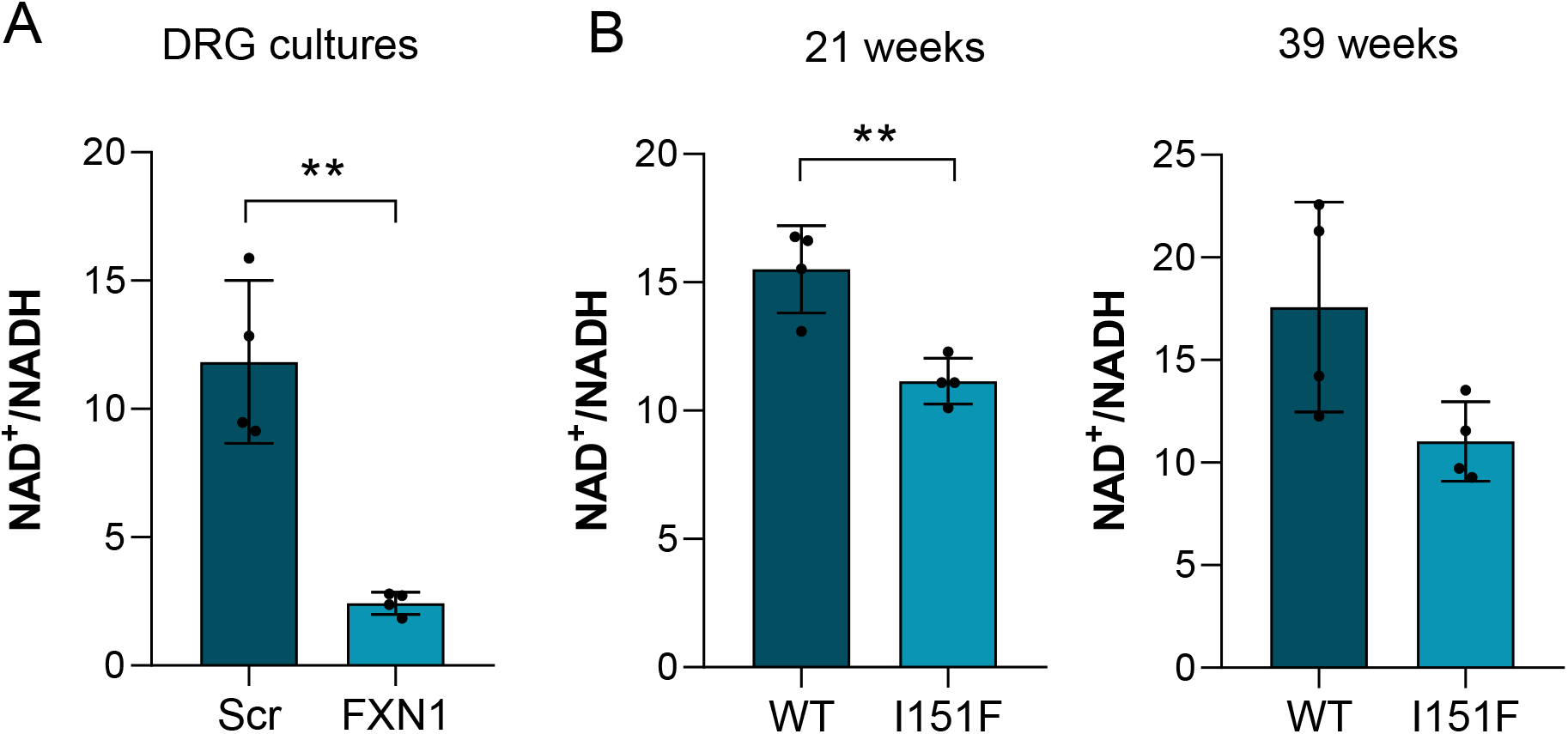
Reduced levels of NAD^+^/NADH ratio in primary cultures of DRG neurons and DRGs from FXN^I151F^ mice. (A) NAD^+^/NADH ratio was measured in frataxin-deficient DRG neurons (FXN1) compared to control cells (Src) at day 5 after lentivirus transduction (n=3 independent cultures). (B) NAD^+^/NADH ratio measured in DRG homogenates from 21- and 39-week-old FXN^I151F^ mice and compared to WT mice (n=3 mice/group). Data represent the mean ± SD. Significant differences are indicated (p values < 0.05(*), 0.01(**), or 0.001(***)).

It has previously been reported that increased NAD^+^ levels in a FA cardiomyopathy model improved cardiac function in a SirT3-dependent manner [39]. In mitochondria, SirT3 is the main NAD^+^-dependent deacetylase, and SirT3 inhibition and protein hyperacetylation have a negative feedback effect contributing to cardiomyopathy in FA [25] [40]. Although the role of SirT3 in FA cardiomyopathy is known and several acetylated proteins have been described [40], there is no information on how NAD^+^ reduction may affect protein acetylation in sensory neurons. We determined levels of SirT3 and sirtuin activity in our two models. Western blotting revealed a mild but significant decrease (15%) in SirT3 levels in FXN1 and FXN2 cells compared to Scr cells (Fig. 5A and B). In DRG homogenates from 21- and 39-week-old FXN^I151F^ mice, SirT3 levels were reduced by around 25%, compared to their controls (although at 39 weeks the difference was not statistically significant) (Fig. 5C and D).

**Fig. 5.**
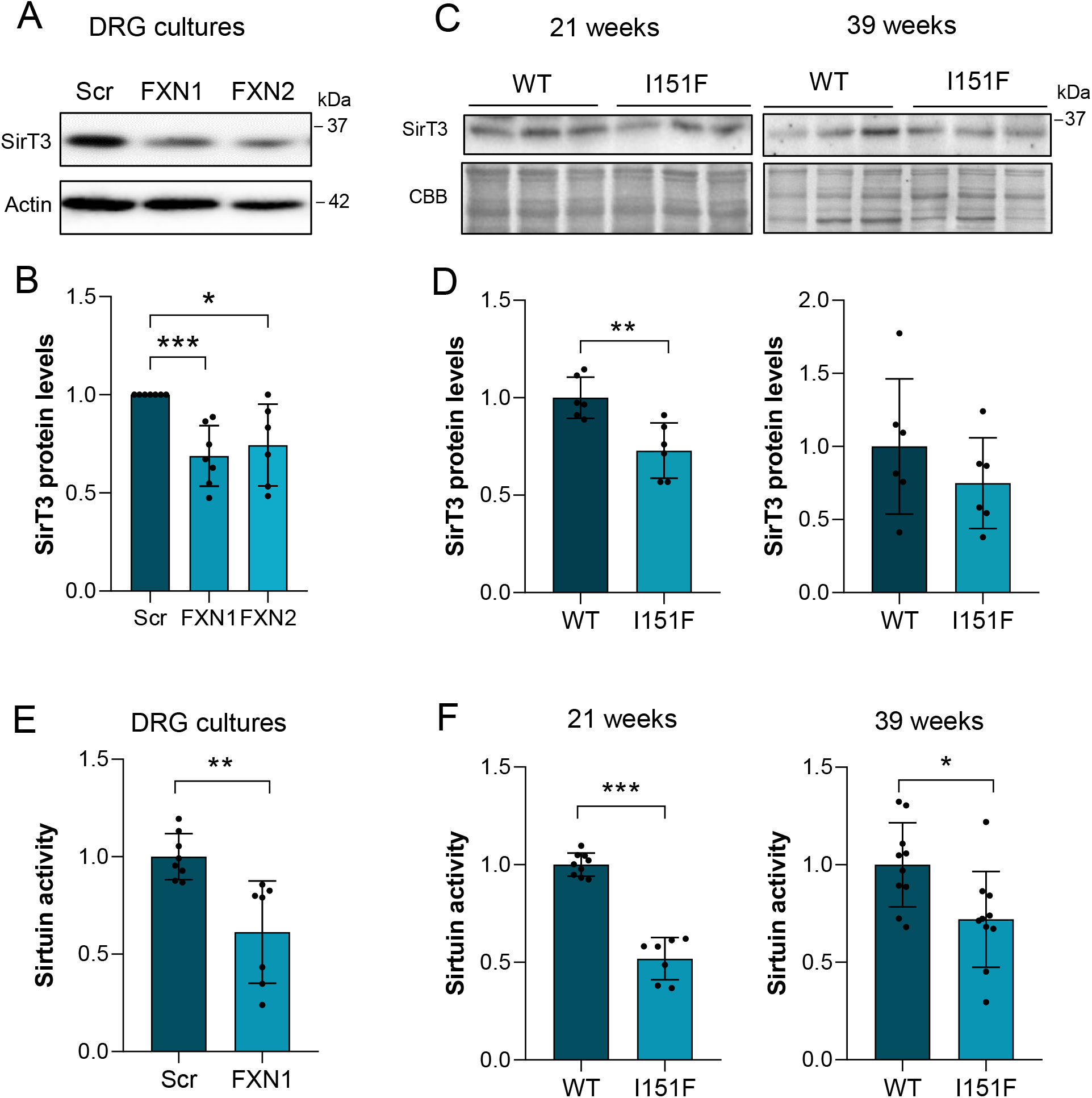
Decreased Sirt3 protein levels and sirtuin activity in frataxin-deficient DRG neurons and DRG homogenates from the FXN^I151F^ mice. (A) Western blot analysis of SirT3 protein levels from frataxin-deficient DRG neurons (FXN1) compared to control cells (Src) at day 5 after lentivirus transduction. Actin was used as a loading control. (B) Quantification data of A are mean ± SD obtained from n=7 independent cultures. (C) SirT3 protein levels were analyzed in DRG homogenates from 21- and 39-week-old FXN^I151F^ mice and compared to WT mice. (D) Quantification data of C are mean ± SD obtained from n=6 mice/group. Coomassie Brilliant Blue (CBB) protein stain was used as a loading control. (E) Sirtuin activity in DRG homogenates from frataxin-deficient DRG neurons (FXN1) compared to control cells (Src) was measured at day 5 after lentivirus transduction. Trichostatin A was added to the assay to inhibit NAD-independent histone deacetylases. Quantification data are mean ± SD obtained from n=3 independent isolations. (F) Sirtuin activity was measured in DRG homogenates from 21- and 39-week-old FXN^I151F^ mice and compared to WT mice. Quantification data are mean ± SD obtained from n=3 mice/group. Significant differences are indicated (p values < 0.05(*), 0.01(**), or 0.001(***)).

Both NAD^+^/NADH and SirT3 reduction should affect SirT3 activity. Since it was impractical to analyze only SirT3 activity in our primary cultures, total sirtuin activity was measured, with SirT1, SirT2 and SirT3 being the sirtuins with the highest deacetylase activity [20]. The results showed a 35–40% decrease in sirtuin activity in FXN1 cells compared to Scr (Fig. 5E). Sirtuin activity was also analyzed in DRG homogenates from FXN^I151F^ and WT mice. As shown in Fig. 5F, a 40% (21-week-old) and 30% (39-week-old) reduction was detected in the mutant mice compared to WT mice.

### 3.4. Increased protein acetylation in FXN-deficient DRGs

A reduction in sirtuin activity would be expected to lead to an increase in protein acetylation. To that end, western blots anti acetyl-Lys were performed using DRG homogenates from WT and mutant mice at 21 and 39 weeks of age. A highly prominent acetylated band of 52 kDa appeared in all samples (Fig. 6A). At 21 weeks, this band presented a mild increase in acetylation in the mutant mice compared to WT animals (not significant). However, at 39 weeks, the differences in protein acetylation between WT and mutant mice increased by up to 2.5-fold (Fig. 6B). We identified this protein as alpha tubulin using 2-dimensional gel electrophoresis followed by mass spectrometry (Supplementary Fig. 4A and B). Alpha-tubulin is regulated by acetylation, among several other posttranslational modifications. Since specific antibodies for acetylated alpha tubulin at the Lys40 residue were available, western blot was performed (Fig. 6C). Similarly to that observed with the acetyl-Lys antibody, we corroborated that tubulin acetylation was significantly increased in 39-week-old FXN^I151F^ mice compared to WT mice (Fig. 6D). At 21 weeks of age, there was a tendency for acetylation to increase, but the increase was not significant.

**Fig. 6.**
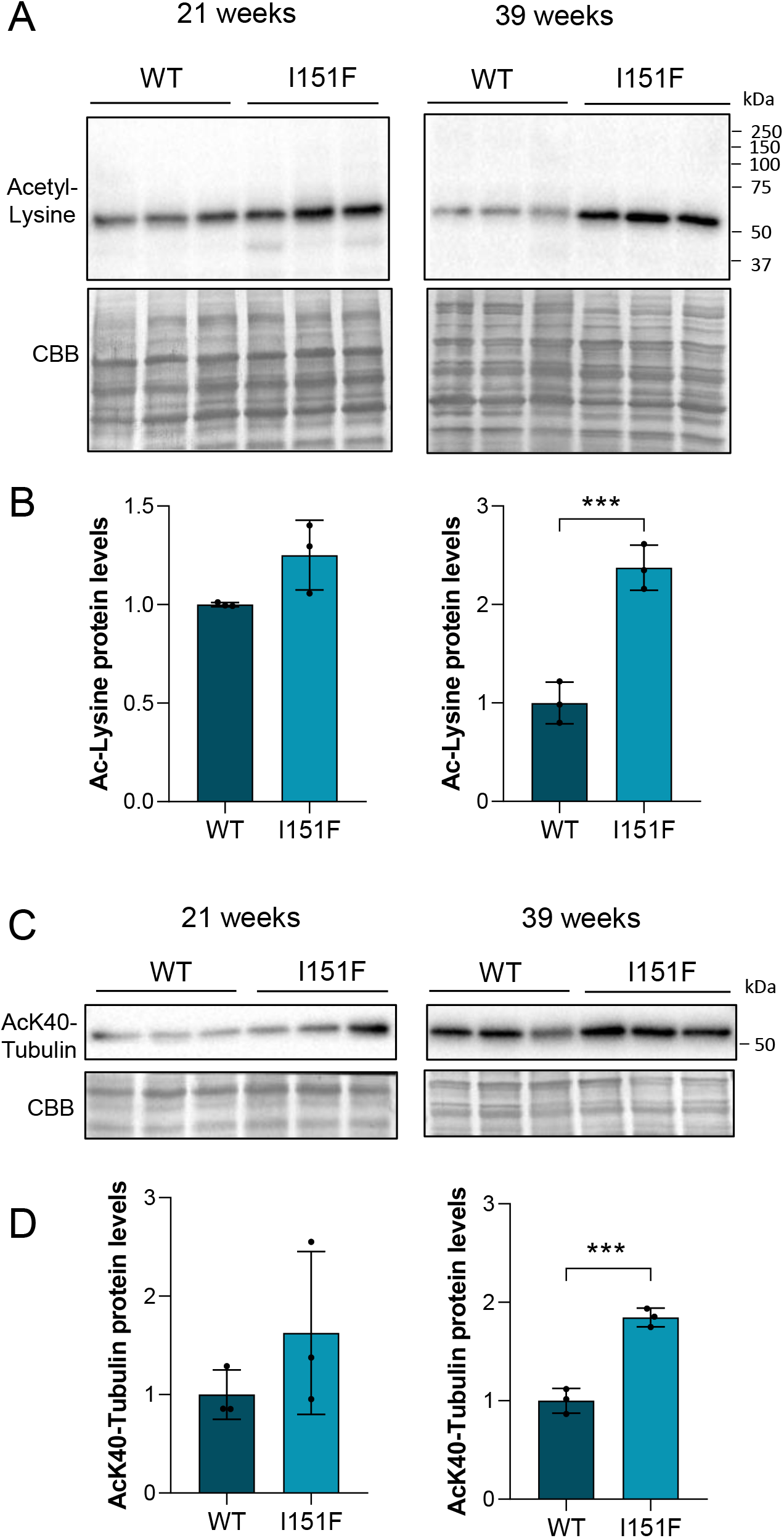
Identification and analysis of alpha tubulin as a highly acetylated protein. (A) Western blot analysis of acetylated proteins from DRG homogenates from 21- and 39-week-old WT and FXN^I151F^ mice. (B) Quantification data of A are mean ± SD obtained from n=3 mice/group. Coomassie Brilliant Blue (CBB) protein stain was used as a loading control. (C) Western blot of alpha-tubulin acetylated at Lys40 from DRG homogenates from 21- and 39-week-old WT and FXN^I151F^ mice. (D) Histogram data from C are represented as means ± SD from n=3 mice/group. CBB protein stain was used as a loading control. Significant differences are indicated (p values < 0.05(*), 0.01(**), or 0.001(***)).

### 3.5. SOD2 acetylation and oxidative stress increase in frataxin-deficient DRGs

A well-known target of SirT3 is mitochondrial SOD2. To understand how decreased sirtuin activity and SirT3 levels may affect it, acetylated SOD2 and total SOD2 were analyzed in 21- and 39-week-old FXN^I151F^ and WT mice (Fig. 7A). As shown in Fig. 7B, the results demonstrated increased acetylation at both ages (21 and 39 weeks) in the mutant mice. Intriguingly, at 21 weeks, the difference in SOD2 acetylation between mutant and WT mice was much higher than at 39 weeks, once normalized by total SOD2 levels (Fig. 7B and Supplementary Fig. 5). It is well known that acetylation at Lys68 inactivates SOD2 [41], and we hypothesize that this might result in oxidative stress. To that end, hydroxynonenal (HNE), a well-studied oxidative stress marker, was analyzed (Fig. 7C). HNE is an α,β-unsaturated hydroxyalkenal generated as a byproduct of lipid peroxidation that can be detected and quantified by western blot, because it covalently binds to specific proteins. As shown in Fig. 7D, HNE was increased in DRG homogenates from FXN^I151F^ mice both at 21 and 39 weeks of age compared to WT mice.

**Fig. 7.**
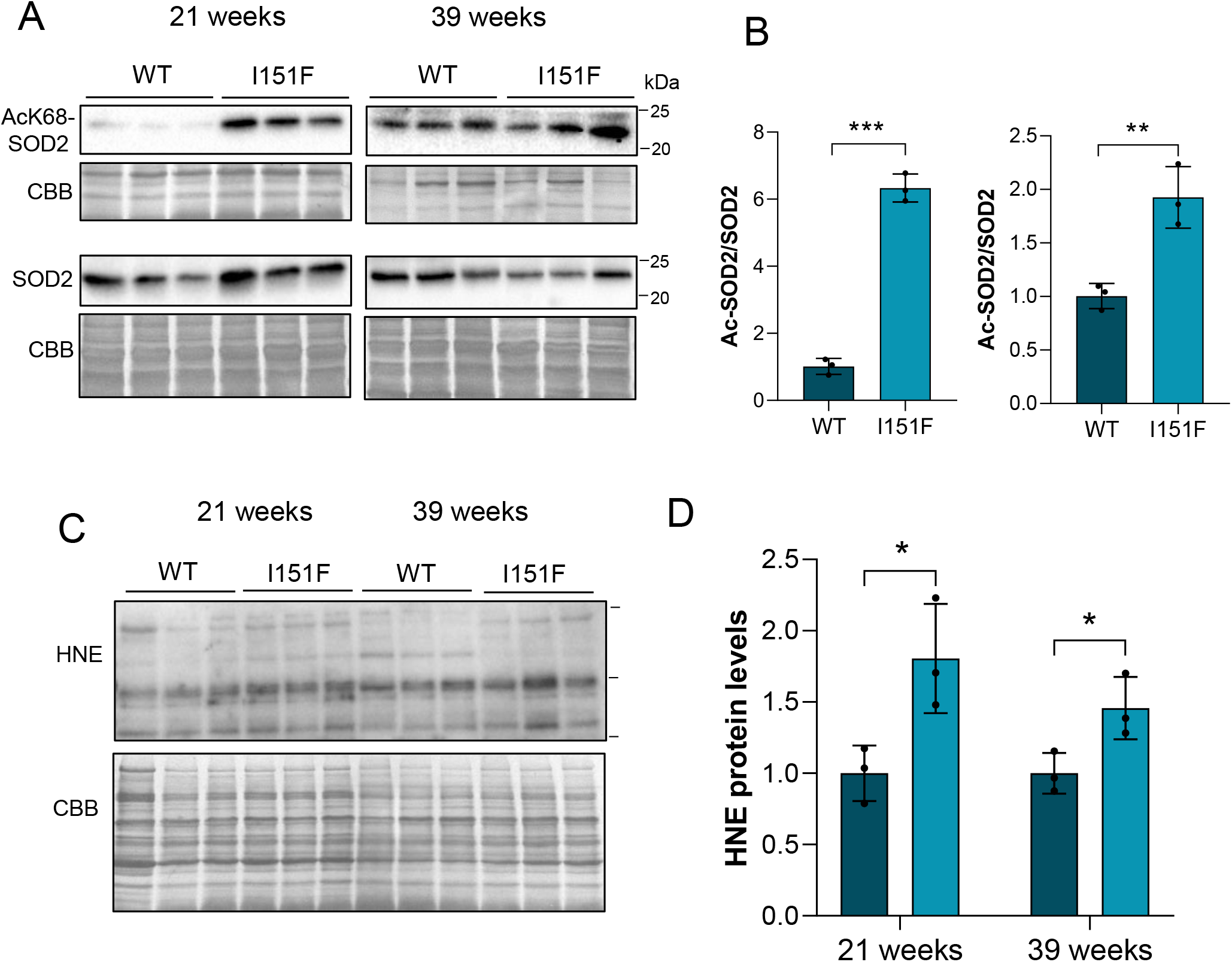
Increased acetylated SOD2 and HNE levels in the mutant mice. (A) SOD2 acetylated at Lys68 and total SOD2 were analyzed by western blot of DRG homogenates from 21- and 39-week-old WT and FXN^I151F^ mice. (B) Histograms represent the mean ± SD from n=3 mice/group. CBB stain was used as a loading control. (C) HNE bound to proteins was analyzed by western blot from DRG homogenates of 21- and 39-week-old mice. D) Histograms represent the mean ± SD from n=3 mice/group. CBB stain was used as a loading control. Significant differences between WT and mutant animals are indicated (p values < 0.05(*), 0.01(**), or 0.001(***)).

To study whether such oxidative stress could also be observed in primary cultures of DRG neurons, and because SOD2 plays such an important role inside the mitochondria, we analyzed the acetylation of this antioxidant enzyme in Scr, FXN1 and FXN2 cultures (Fig. 8A). The results showed that FXN1 and FXN2 cells presented a 2- to 3-fold increase in SOD2 acetyl-Lys68 levels compared to Scr, once normalized by total SOD2 levels (Fig. 8B and Supplementary Fig. 5). To study the effect of increased SOD2 acetylation on frataxin-deficient DRG neurons, mitochondrial superoxide levels were analyzed using the fluorescent dye MitoSox Red (Fig. 8C). As shown in Fig. 8D, FXN2 cultures with reduced frataxin levels showed increased mitochondrial superoxide levels compared to Scr cells.

**Fig. 8.**
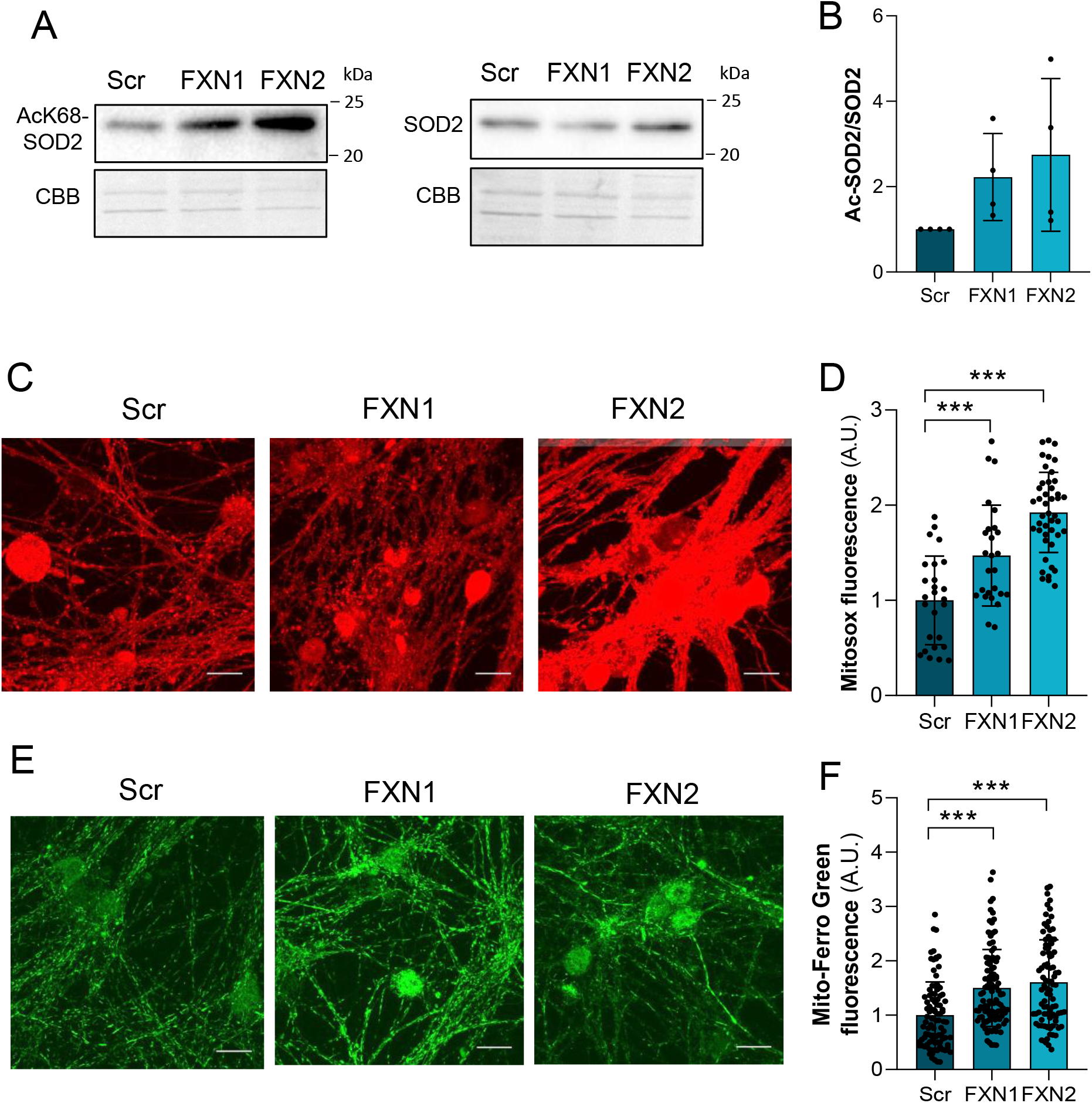
Oxidative stress increases in frataxin-deficient DRG primary cultures. (A) SOD2 acetylated at Lys68 and total SOD2 were analyzed by western blot in Scr (control), FXN1 and FXN2 (5 days after lentivirus transduction) primary culture homogenates. Representative western blot images are shown. (B) Histograms represent the mean ± SD from 4 independent isolations. CBB: Coomassie Brilliant Blue stain was used as a loading control. (C) Mitochondrial superoxide levels were analyzed using the fluorescent dye MitoSox Red in frataxin-deficient DRG neurons (FXN1 and FXN2) and compared to control cells (Src) at day 5 after lentivirus transduction. (D) Histograms represent fluorescence intensity in soma and data are mean ± SD. Between 17 and 35 fields were analyzed for Scr conditions, between 20 and 40 fields for FXN1 and between 19 and 37 fields for FXN2. For each condition, three images per field were taken. (E) Mitochondrial Fe^2+^ staining with Mito-Ferro Green probe in primary cultures of DRG neurons at day 3 after lentivirus transduction. Scale Bar = 100 μm and objective = 100x with confocal microscope Olympus FV1000. (F) Data are mean ± SD obtained from n=3 independent isolations. Between 88 and 104 soma were analyzed for Scr, FXN1 and FXN2. In D and F, significant differences between Scr and FXN1 or FXN2 are indicated (p values < 0.05(*), 0.01(**), or 0.001(***)).

Another trait that can cause oxidative stress is iron accumulation, especially ferrous iron (Fe^2+^). Published evidence indicates that frataxin deficiency causes an imbalance in iron homeostasis. Iron deposits or accumulation have been observed in frataxin-deficient models [42], but their role in the progression of the disease remains unclear [43]. To analyze whether iron accumulates in sensory neurons of DRG, the fluorescent probe Mito-Ferro Green was used (Fig. 8E). This compound enables live cell fluorescent imaging of intramitochondrial Fe^2+^. Quantification of mito-Ferro Green fluorescence in the soma of DRG neurons showed around 1.5–2-fold iron accumulation in FXN1 and FXN2 cells compared to Scr cells (Fig. 8F). In summary, all these changes contribute to oxidative stress that affects the whole cell but in particular the mitochondria in the sensory neurons.

### 3.6. Honokiol alleviates mitochondrial dysfunction

To better understand the role of SirT3 in the mitochondrial deterioration observed in frataxin-deficient cells, Scr, FXN1 and FXN2 primary cultures were treated with honokiol, a well-known SirT3 activator [27]. As shown in Fig. 9, honokiol treatment reduced the decay in oxygen consumption observed in FXN1 and FXN2 compared to Scr cells. The recovery observed was significant under all conditions analyzed, but was higher in FXN1 and FXN2 than in Scr cultures (Fig. 9B - D).

**Fig. 9.**
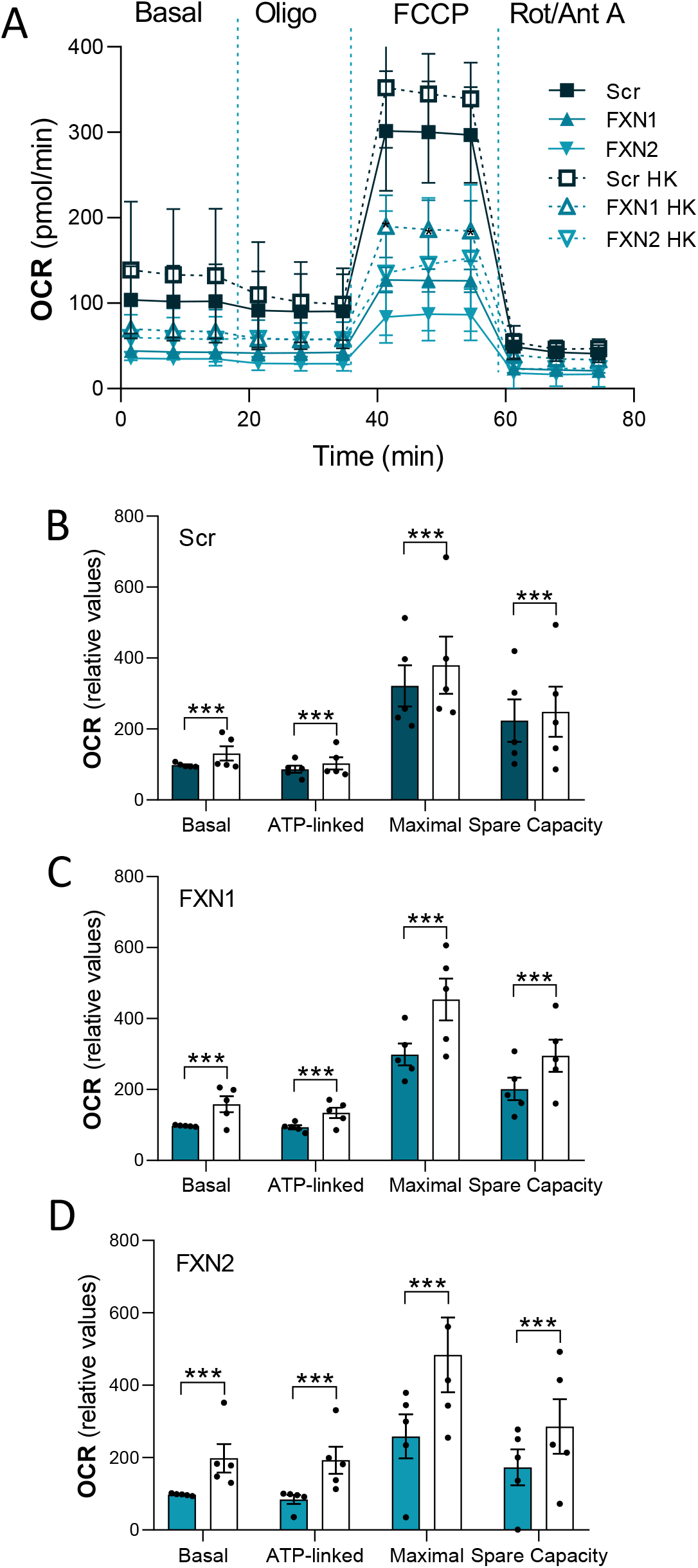
Honokiol treatment ameliorated the impairment in oxygen consumption detected in frataxin-deficient primary cultures. (A) Mitochondrial stress analysis of frataxin-deficient DRG neurons (FXN1 and FXN2) compared to control cells (Src) at day 5 after lentivirus transduction (n=4 independent cultures). Honokiol (2 µM) was added simultaneously with the virus (day 0) and 3 days later. (B-D) Basal respiration, ATP production, maximal respiration rate, and spare capacity of Src cultures (B) compared to FXN1 (C) and FXN2 (D). Values are indicated as relative to basal respiration, and are mean ± SD obtained from n=4 independent isolations.

## 4. Discussion

Degeneration of DRG large sensory neurons is one of the initial events in FA. However, the effect of frataxin deficiency on their mitochondrial metabolism has not been properly studied [38]. In the present study, we demonstrate that mitochondrial metabolism is impaired in both primary cultures of frataxin-deficient DRG neurons and DRGs from the FXN^I151F^ mouse. In the primary culture model, decreasing levels of frataxin correlated with increasing reductions in oxygen consumption. After 5 days of lentivirus transduction, when frataxin levels were similar to those found in human patients, cells exhibited a strong and consistent decrease in both the OCR (basal and maximal) and total ATP production. As expected, mitochondrial ATP production was strongly impaired but a mild decrease in glycolytic ATP production was also observed. Consequently, mitochondrial membrane potential decreased, as previously described [11].

To determine whether these effects also occur *in vivo*, we analyzed the consequences of the FXN^I151F^ mutation in mice, which is equivalent to the human I154F pathological mutation. These mice present systemic frataxin deficiency and severe phenotypes, such as decreased weight gain (which is observed from 15 weeks onward), neurological deficits (which start at 23 weeks) and marked biochemical alterations, which become apparent before the appearance of the functional alterations [29]. Based on this information, we conducted our study in mice at 21 weeks of age, before neurologic symptoms appear, and at 39 weeks of age, when the neurological symptoms are clearly apparent. DRGs were isolated from mutant and WT mice and the homogenates were analyzed. Among the complexes of the OXPHOS system, complexes I and II were the most affected. We found a reduction in the content of NDUFB8 (complex I) and SDHB (complex II), consistent with a decline in their enzyme activity. These results are also consistent with the lower levels of complexes I and II observed in the cerebrum, cerebellum, and heart of FXN^I151F^ mice [29]. They also concur with previous results in the KIKO mouse model, where these two complexes showed decreased enzyme activity in cerebella from asymptomatic mice [44]. In the cerebellum, gene expression analysis indicated that this decrease was due to posttranscriptional events [29]. Since NDUF8 and SDHB contain iron-sulfur clusters or belong to complexes containing them, it may be hypothesized that these proteins are degraded due to the absence of cofactor. However, we should be cautious, since FXN^I151F^ mice do not have a general loss of iron-sulfur clusters [29]. Therefore, the mechanism explaining the loss of these proteins may be more complex and could involve different regulatory pathways. It has been reported that astrocyte reactivity contributes to the progression of the disease and that neuron– glia interactions are also important to prevent neurodegeneration in FA [45].

Mitochondrial impairment results in a decreased NAD^+^/NADH ratio in primary cultures of frataxin-deficient DRG neurons as well as in DRGs from FXN^I151F^ mice. Accordingly, as an NAD^+^-dependent enzyme, mitochondrial SirT3 activity is expected to decrease in both models. The role of SirT3 in cardiac metabolism has been well described [19] [46] In the context of FA, SirT3 has been studied in two conditional mouse models that develop a fatal cardiomyopathy and impaired activity in respiratory complexes [40]. They showed marked hyperacetylation of cardiac mitochondrial proteins and a huge decrease (85-fold) in the NAD^+^/NADH ratio. It has also been reported that, using the cardiac/skeletal muscle–specific FXN-KO, activation of SirT3 by nicotinamide mononucleotide supplementation improves both cardiac and extracardiac metabolic function and energy metabolism [19]. In addition, two subunits of complex II, SDHA and SDHB, interacted specifically with SirT3 and were deacetylated and activated by this mitochondrial sirtuin [22].

However, the role of SirT3 in the nervous system and, especially, in sensory neurons has not been studied in FA. In the present study, we observed a 5-fold decrease in the NAD^+^/NADH ratio in FXN1 cell cultures compared to Scr. In FXN^I151F^ mice, the ratio decreased by 30–40% compared to WT animals, a mild reduction compared to results published by Wagner et al. [40] using mitochondria isolated from heart. Such differences may be attributed to the fact that we measured the NAD^+^/NADH ratio in total DRG homogenates, not mitochondrial preparations. In the present study, the small size of the DRGs did not allow us to purify enough mitochondria to perform this analysis. In addition, DRGs and heart may have different NAD^+^/NADH ratios, and Wagner et al. used a mouse model that has a complete deficiency of frataxin.

Nevertheless, our NAD^+^/NADH data are consistent with the reduced sirtuin activity observed. Since whole homogenates were used, we measured total sirtuin activity (NAD^+^-independent deacetylase activities were fully inhibited). Among the seven sirtuins present in mammalian cells, extramitochondrial SirT1 and SirT2, and predominantly mitochondrial SirT3, showed the highest deacetylase activity [47], while other sirtuins performed deacylation reactions other than acetyl groups (malonyl, succinyl, palmitoyl,…) and the ADP-ribosylation reaction [20]. Three sirtuins, SirT3, 4 and 5, are located predominantly within the mitochondrial matrix. SirT3 shows robust deacetylase activity while that of SirT4 and 5 is weak or null [19] [20]. The results presented here showed a 40% decrease in sirtuin activity in FXN1 cultures compared to Scr. This decrease was around 35–50% in total DRG homogenates from FXN^I151F^ mice compared to WT animals. These results indicate that: i) it is highly possible that in mitochondria the differences in SirT3 activity between mutant and WT animals could be higher; and ii) the impairment in the mitochondrial metabolism caused by frataxin deficiency probably disturbs the entire cell metabolism.

It has been reported that oxidative stress induces sirtuin levels [48] [49]. The SirT3 promoter has tandem Nrf2 consensus binding motifs [50] suggesting that Nrf2 regulates *SIRT3* expression and may explain *SIRT3* upregulation in response to various stressors. However, in the FA context, previously published results showed no differences in SirT3 protein levels in NSE-KO heart [40] or FXN-KO heart [39]. We observed moderate but significantly decreased SirT3 levels, both in FXN1 and FXN2 primary cultures, and DRGs from FXN^I151F^ mice (at 21 and 39 weeks of age) compared to controls. This decline may be due to i) decreased synthesis, since reduced SirT3 mRNA expression was detected in FXN-KO heart [39]; or ii) increased protein degradation after oxidative modification. In fact, SirT3 modification by 4-HNE was much higher in heart from frataxin-deficient mice compared to WT mice [40]. Such carbonyl group adduction has also been described in other pathological conditions, for example, in alcoholic liver disease [51], where the covalent modification of SirT3 by 4-HNE at Cys280 (a critical zinc-binding residue) was identified by tandem mass spectrometry. This modification resulted in the allosteric inhibition of SirT3 activity [51].

As already indicated, most of the available knowledge on SirT3 comes from studies in non-neuronal cells. Targets of SirT3 deacetylation have been analyzed in the heart [19], but the interactions between SirT3 and its protein substrates in the CNS are only just starting to be elucidated [47]. In DRGs from FA models, no information was available. In this work, acetylated proteins analyzed by western blot after one-dimensional SDS-PAGE showed a pattern with a predominant 52 kDa acetylated protein. This protein was identified by mass spectrometry after separation by 2D electrophoresis as alpha tubulin. Western blot anti acetyl-tubulin demonstrated increased alpha tubulin acetylation in mutant mice at 21 weeks, but more clearly at 39 weeks of age, compared to WT mice. Tubulin, composed of heterodimers of alpha and beta tubulin, is the main component of microtubules, which play important roles in cell motility, mitosis, and intracellular vesicle transport. Tubulins are highly structured proteins, and both alpha and beta tubulin undergo many posttranslational modifications, including acetylation/deacetylation [52]. Neuronal alpha-tubulin acetylation occurs at Lys40 in the N-terminal region [53]. The histone deacetylase HDAC6 and SirT2 have been identified as alpha-tubulin deacetylases [52]. Reversible acetylation of alpha-tubulin may be involved in regulating microtubule stability, cell motility, and axon regeneration. However, the role of alpha-tubulin acetylation in neurons is very complex and not fully understood. On the one hand, it has been reported that Lys40 acetylation reduces the flexural rigidity of microfilaments, protecting them from damage after repetitive bending [54]. On the other hand, several reports have linked alpha-tubulin acetylation to neurodegeneration [55] [56].

SirT3 activates the mitochondrial manganese-dependent SOD2 through deacetylation at the Lys68 site [57]. SOD2 is an essential antioxidant enzyme against the superoxide radicals generated inside mitochondria, mainly by the ETC. It has been reported to protect neurons against degeneration in several models of neurodegenerative disorders [58] [59]. We showed increased SOD2 acetylation at Lys68 in DRGs from the FXN^I151F^ mice. Interestingly, when compared to WT animals, a 6-fold increase in acetylation was detected at 21 weeks of age while at 39 weeks, there was only a 2-fold increase. The opposite pattern was observed for alpha-tubulin, where the difference in acetylation versus the WT animals was higher at 39 weeks. This may indicate that mitochondrial SirT3 substrates are early targets while cytosolic proteins are acetylated later when the neurologic symptoms are evident. This boost in acetyl-SOD2 at 21 weeks would generate oxidative stress that would be more apparent at this initial stage of the disease. The amount of HNE bound to proteins is consistent with this idea, since levels in the mutant mice were higher at 21 weeks compared to 39 weeks. In agreement with our results, it has been reported that the SOD2 mutant mice (CD1-Sod2^tm1Cje^) carrying inactive SOD2 exhibit inhibition of the respiratory chain complex I (NADH-dehydrogenase) and complex II (succinate dehydrogenase) [60].

By deacetylating SOD2, SirT3 protects neurons against metabolic and oxidative stress. It achieves this by reducing mitochondrial superoxide levels, stabilizing cellular and mitochondrial Ca^2+^ homeostasis, and inhibiting mitochondrial membrane transition pore opening to prevent apoptosis [61]. In primary cultures, we showed increased acetylation of SOD2 at Lys68 in FXN1 and FXN2 cells compared to Scr and a correlation with higher levels of superoxide anion in frataxin-deficient cells. In fact, previous results from our group also showed Ca^2+^ dyshomeostasis and MPTP opening in FXN1 and FXN2 primary cultures [11].

Honokiol (HKL) is a biphenolic compound derived from the bark of magnolia trees and a widely known SirT3 activators. It has anti-inflammatory, anti-tumor, anti-oxidative, and neuroprotective properties [27]. Honokiol has shown some therapeutic effects in heart diseases, cancer, and metabolic diseases [62] but the mechanism(s) remains unclear. Honokiol enhances SirT3 expression and can be found in mitochondria [62] and a non-allosteric mechanism of activation of Sirt3 has also been reported [63]. In the nervous system, honokiol has been reported to attenuate oxidative stress and neuroinflammation in the hippocampus [64]. It also mitigates oxidative stress and mitochondrial dysfunction by regulating mtROS homeostasis, partly via the AMPK/PGC-1α/Sirt3 pathway [65]. In Alzheimer disease, honokiol increased SirT3 expression levels and activity, which in turn markedly improved ATP production and weakened mitochondrial ROS production, rescuing memory deficits [66]. When the compound was added to our primary cultures, an improvement in oxygen consumption was observed, confirming the role of SirT3 in the mitochondrial pathology caused by frataxin deficiency. It should be noted that the recovery was partial, indicating that perhaps honokiol was not able to fully recover SirT3 activity to WT levels. Other pathological alterations may also be relevant in mitochondrial impairment, including a deficit in Fe-S cluster biosynthesis and iron accumulation.

Although it remains a matter of debate, deficiency of Fe-S clusters could be a secondary trait in frataxin-deficient cells because FXN^I151F^ mice do not present a general loss of Fe-S clusters [29], and similar results were observed in the yeast model of FA [67]. Iron is the most abundant transition metal element within an organism, but because of its high reactivity, free iron and especially the reduced form ferrous ion, can catalyze the Fenton reaction, generating the highly reactive hydroxyl radical. Using the probe Mito-Ferro Green we demonstrated that frataxin-deficient DRG primary cultures accumulated mitochondrial Fe^2+^ compared to control cultures. A recent report described several conserved amino acid residues in the frataxin proteins, grouped in four clusters. It was hypothesized that cluster 3, present only in eukaryotic and Rickettsia frataxins but not in their bacterial homologs, may help to prevent the formation of ROS during iron detoxification [68]. Altogether this raises the possibility that ferroptosis may play a role in cellular death in sensory neurons, as has been described in several models of FA [13] [14]. Although the increase in HNE-protein adducts suggests such a possibility, this hypothesis will need to be explored further.

## 5. Conclusions

The lack of frataxin induces, probably in multiple ways, a decrease in ETC and ATP synthesis. Such mitochondrial dysfunction reduced the NAD^+^/NADH ratio leading to lower sirtuin activity. We have provided evidence that the decrease in SirT3, a key regulator of mitochondrial energetic and antioxidant metabolism, resulted in oxidative stress due to increased SOD2 acetylation, which inactivates the enzyme. As a consequence, higher levels of superoxide in the mitochondria were detected, as well as increased HNE (a lipid peroxidation byproduct). In parallel with the frataxin deficiency, this may explain the Fe^2+^ accumulation, which exacerbates the oxidative stress. We propose that because neurons have a high energy requirement, and fewer antioxidant systems compared to other tissues, the decay in the ETC results in decreased SirT3 activity and protein hyperacetylation, thus comprising a negative feedback contributing to neuron lethality.

## Abbreviations

DRG: Dorsal root ganglia
ECAR: extracellular acidification rate
ETC: electron transport chain
FA: Friedreich ataxia
HNE: 4-Hydroxynonenal
OCR: Oxygen consumption rate
SOD2: Mn-superoxide dismutase
ROS: reactive oxygen specie

## Ethics statement

Animal use and care protocol conforms to the National Guidelines for the regulation of the use of experimental laboratory animals from the Generalitat de Catalunya and the Government of Spain (article 33.a 214/1997) and was evaluated and approved by the Experimental Animal Ethical Committee of the University of Lleida (CEEA).

## Declarations of interest

The authors declare no competing interests.

## Acknowledgments

We thank Roser Pané for her excellent technical assistance.

## Funding

This work was supported by Ministerio de Economía y Competitividad, MINECO (Spain) [grants PN-P21018 and PDC-N21019] and Generalitat de Catalunya [SGR2009-00196]. ASA received a Ph.D. fellowship from the Generalitat de Catalunya. Maria Pazos received a PhD fellowship from the Universitat de Lleida.

## Author contributions

Conceptualization, design and perform experiments: A.S-A., F.D., J.T., M.M-C, M.P-G and E.C. Data analysis: A.S-A, JR, and E.B. Histochemical examinations: A.S-A, M.P-C. Primary cell cultures and mice control: A.S-A, M.P-C., and F.D. Writing original draft: A.S-A, J.R. and E.C. Reviewed and editing: J.T., F.D., J.R. and E.C.

All authors have read and agreed to the published version of the manuscript.

## Data and materials availability

All data required to evaluate the conclusions in the paper are present in paper and the supplementary materials.

## Notes

### Competing Interest Statement

The authors have declared no competing interest.

