## Supplemental figures for "Deficient mitochondrial respiration impairs sirtuin activity in dorsal root ganglia in Friedreich Ataxia mouse and cell models"

Supplementary figure 1

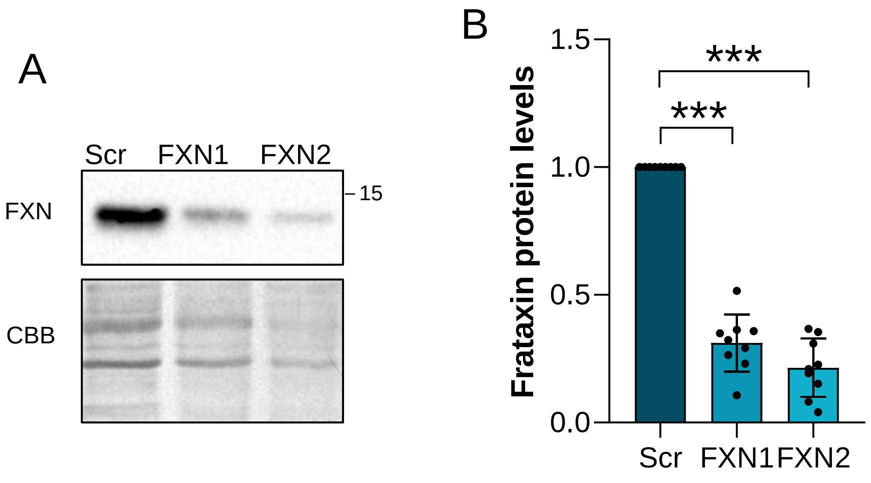


**Frataxin decreases after lentivirus transduction**
(A) FXN amounts were analyzed by Western blot in frataxin-deficient DRG neurons at day 5 after lentivirus transduction as described in Materials and Methods. Coomassie Brilliant Blue (CBB) protein stain was used as a loading control. Quantification is shown in (B). Data are mean ± SD obtained from n = 9 independent isolations.

Supplementary figure 2

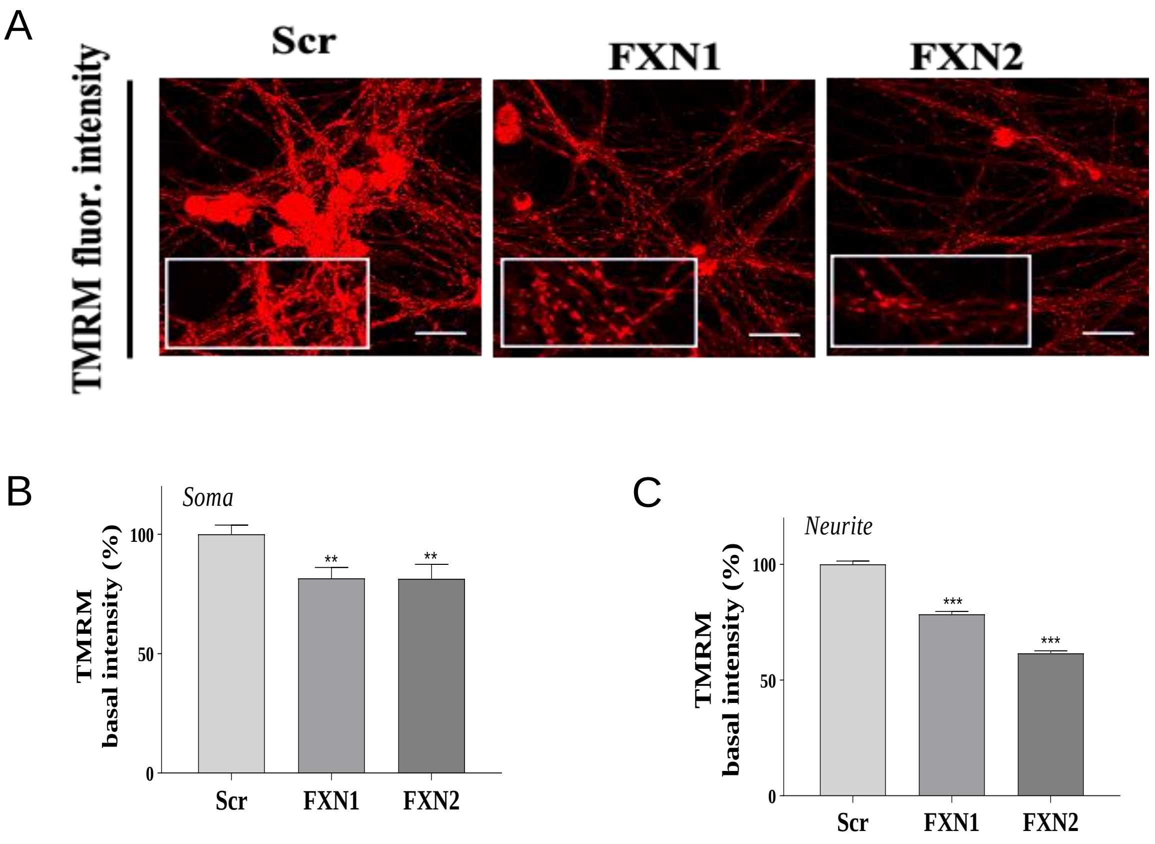


**Frataxin reduction induces mitochondrial depolarization.**

A) Representative images of fluorescent dye TMRM in frataxin deficient DRG neurons at day 5 after lentivirus transduction**.** Scale Bar: 30 µm and × 32 lens.. Histograms in B and C represent fluorescent intensity in soma and neurite respectively. Data are mean **±** SEM obtained from n = 5 independent isolations. A range of 17–35 fields has been analysed for Scr conditions, a range of 20–40 fields for FXN1 conditions and 19–37 fields for FXN2 conditions. For each condition, three images per field have been taken. Significant differences between Scr and FXN1 or FXN2 are indicated (p values < 0.05(*), 0.01(**), or 0.001(***)).

Figure S2A, B and C are preprinted, with permission, from Biochem J. (2021) Jan 15;478(1):1-20. Calcitriol increases frataxin levels and restores mitochondrial function in cell models of Friedreich Ataxia. Britti E, Delaspre F, Sanz-Alcázar A, Medina-Carbonero M, Llovera M, Purroy R, Mincheva-Tasheva S, Tamarit J, Ros J.

Supplementary figure 3

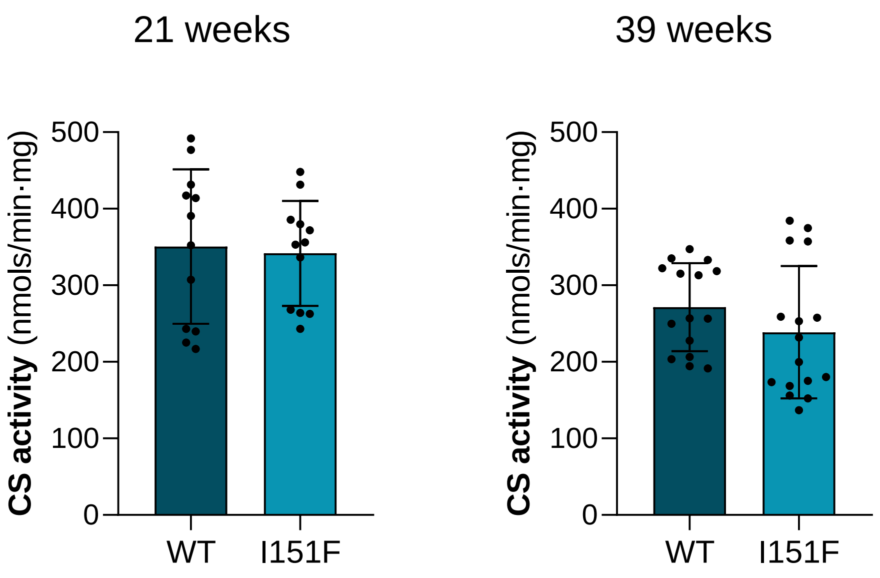


**Citrate synthase activity showed no differences between WT and FXN$^{I151F}$ mice**. Citrate synthase (CS) activity was measured from DRG homogenates from 21- and 39-week-old WT and FXN$^{I151F}$ mice. Data are mean of ± SD from 3 independent experiments. No statistically significant differences between WT and FXN$^{I151F}$ animals were detected either at 21 and at 39-week-old mice.

Supplementary figure 4

A

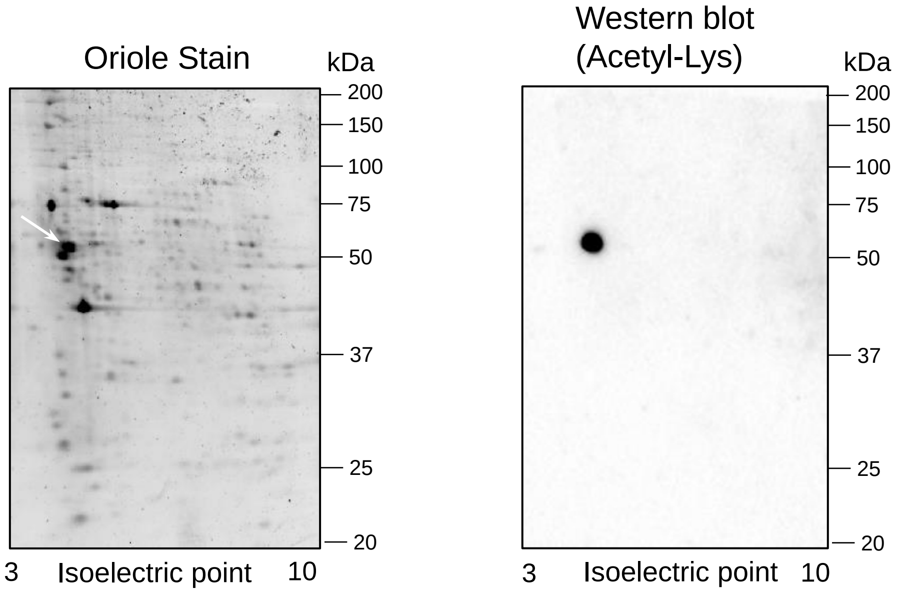


B

| Accession number | Description | Sum PEP Score | Coverage [%] | # Peptides | # PSMs | # Unique Peptides | # AAs | MW [kDa] | Calc. pI | Score Sequest HT: Sequest HT |
|---|---|---|---|---|---|---|---|---|---|---|
| P68369 | Tubulin alpha-1A chain OS=Mus musculus | 82,106 | 58 | 17 | 124 | 6 | 451 | 50,1 | 5,06 | 181,97 |
| P68368 | Tubulin alpha-4A chain OS=Mus musculus | 68,136 | 53 | 16 | 73 | 5 | 448 | 49,9 | 5,06 | 95,07 |
| Q9JJZ2 | Tubulin alpha-8 chain OS=Mus musculus | 47,35 | 33 | 10 | 54 | 1 | 449 | 50 | 5,1 | 69,28 |

PSM: Peptide-spectrum match

**Identification of the highly acetylated protein of DRGs from WT and FXN$^{I151F}$ mice.** (A) Proteins were separated by isoelectric focusing (pH range 3–10, NL) followed by SDS–PAGE and, in parallel, either oriole stain or anti-acetyl Lys Western blot was performed. (B) Acetylated protein was identified by mass spectrometry after in gel tryptic digestion as alpha tubulin. Three isoenzymes (1A chain, 4A chain and 8 chain) were identified with a significant PEP Score.

Acetylation was introduced in the database search analysis queries, but no acetylation was observed. This may probably due because peptide containing Lys40 was not sequenced.

Supplementary figure 5

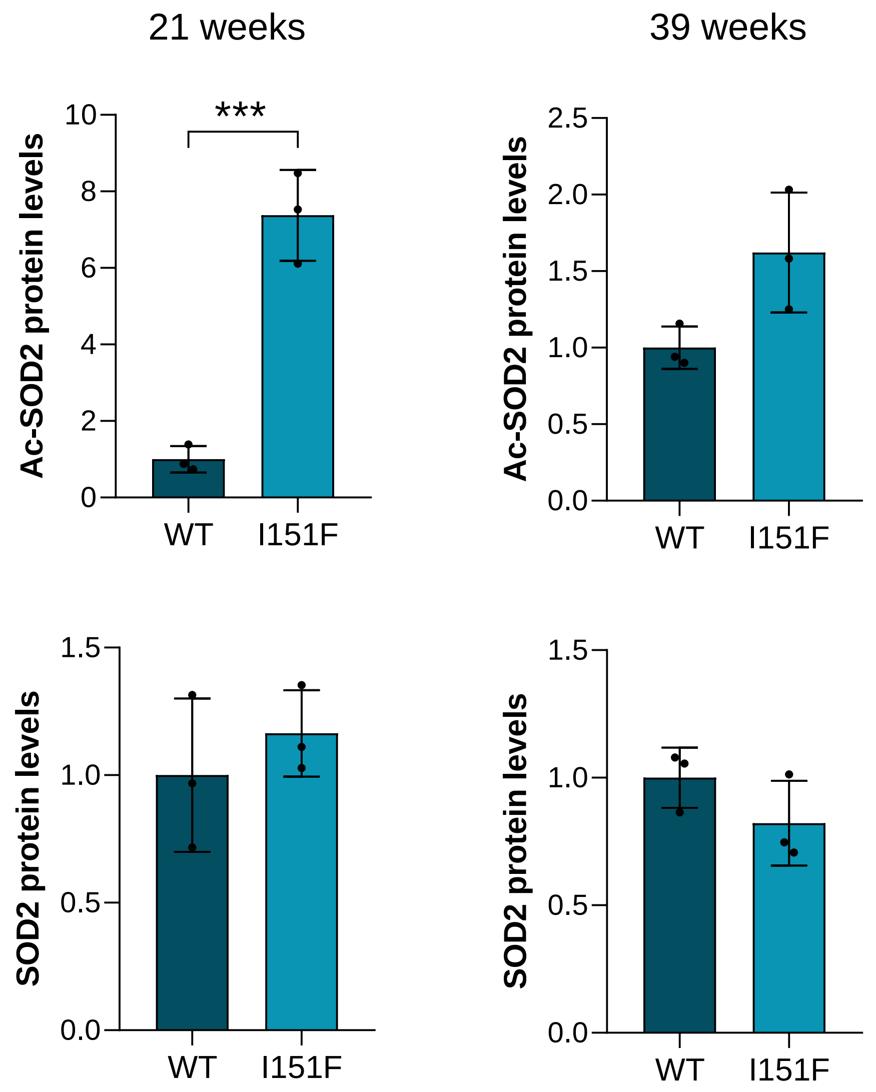


**Sod2 acetylated at Lys68 and total SOD2 levels in DRGs.** DRG homogenates from 21-week and 39-week WT and FXNI151F mice were analyzed by western blot with anti-SOD2 AcK68 and anti-SOD2 antibodies. Coomassie Brilliant Blue stain was used as a loading control. Histograms represent mean ± SD from three mice. Significant differences between WT and mutant animals are indicated (p values < 0.05(*), 0.01(**), or 0.001(***)).

Supplementary figure 6

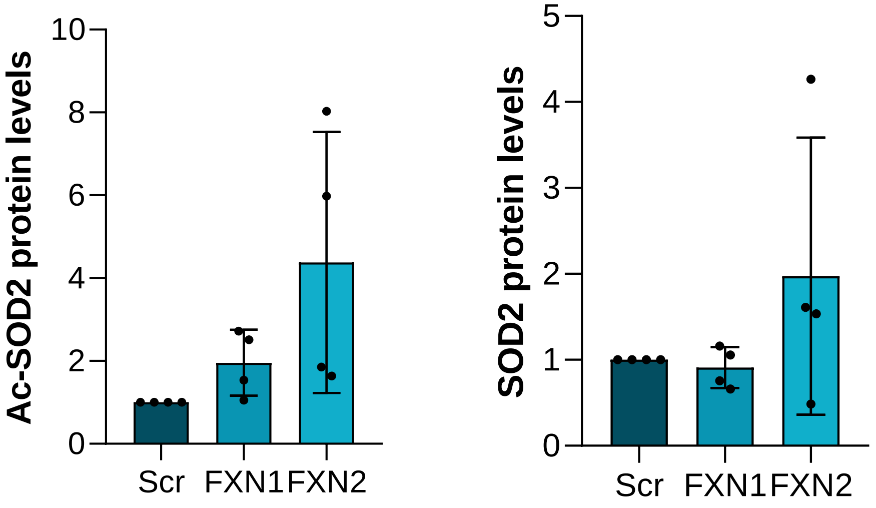


**Sod2 acetylated at Lys68 and total SOD2 were analyzed in DRG neurons**. Scr (control), FXN1 and FXN2 (5 days after lentivirus transduction) primary culture homogenates were analyzed by western blot with anti-SOD2 AcK68 and anti-SOD2 antibodies. Coomassie Brilliant Blue stain was used as a loading control**.** Histograms represent mean ± SD from 4 independent isolations. Significant differences between WT and mutant animals are indicated (p values < 0.05(*), 0.01(**), or 0.001(***)).
